# Steady-state epithelial apical flatness is characterized by MLCK morphodynamics and asynchronous Ca^2+^ oscillations, but not underlying ECM geometry

**DOI:** 10.1101/2025.11.21.688964

**Authors:** Hao Wu, Emerson Herrmann, Jeanette Hyer, Lisa Hua, Takashi Mikawa

**Author notes:** Contributed equally (Hao Wu, Emerson Herrmann). H.W. and T.M. designed the research; H.W. performed research; H.W., E.G.H., J. H., L.L.H., and T.M. analyzed data; and E.G.H., J.H., L.L.H., and T.M. wrote the paper. The authors declare no conflict of interest.

## Abstract

The canonical simple epithelium is a flat sheet-like tissue of horizontally packed cells. While the basal surface is delineated by the basement membrane of extracellular matrix (ECM), little is known about how a flat apical surface is maintained, or if apical/basal dynamics are coordinated. The current study tests the role of the apical domain, to define mechanisms involved in maintaining a flat apical geometry in an epithelium. When the basal geometry is modulated, Madin-Darby Canine Kidney (MDCK) cells adjust their morphology to maintain an overall apical flatness of the confluent layer. Pharmacological and transgenic disruption of non-muscle myosin ATPase, and MLCK activity results in an uneven apical structure, and overall loss of the flat geometry typical of a confluent epithelium. Surprisingly, transgenic experimentation showed that forces maintaining individual MDCK cell flatness are cell-autonomous. Finally, Ca^2+^ imaging reveals an asynchronous calcium flux across a confluent epithelium, suggesting a myosin II-mediated mechanism for maintaining a flat apical architecture. Our results highlight that apical/basal cellular surfaces may not be tightly coordinated, but rather independently regulated. This study provides a new paradigm for how apical flatness is regulated at steady state.

**Impact Statement:**

- How surface flatness of an epithelium is established and maintained is unknown; the data reveal that apical flatness is controlled independently from basement membrane geometry, and require balancing of myosin II morphodynamics.

## Introduction

Simple epithelia sit on a basement membrane of extracellular matrix (ECM) (Martin-Belmonte & Mostov, 2008; Khalilgharibi & Mao, 2021; Kozyrina et al., 2020). At the tissue level, epithelial cells assemble to form a steady-state epithelium, characterized by the balance of cell proliferation and death to maintain a flat, unified apical surface resistant to external stressors (Lemke et al., 2021; Banjac et al., 2023; Odenwald et al., 2018; Tai et al., 2019). Loss of this formation has been implicated in different types of cancer (Hannezo et al., 2014; Eisenhoffer & Rosenblatt, 2012; Peglion & Etienne-Manneville, 2024; Janiszewska et al., 2020; Ribatti et al., 2020; Coradini et al., 2011; Rodriguez-Boulan & Macara, 2014). To date, it is unclear how a uniform flat apical surface is maintained in a steady-state epithelium. In this study, we investigated the underlying mechanism(s) maintaining the flatness of the apical surface, and its relationship with the geometry of the underlying basement membrane in a steady-state epithelium.

An actomyosin network exists at the lumen/cytoplasmic face of the adherens junctions, or apical side, of MDCK epithelial cells (Marivin et al., 2022; Klinger et al., 2014; Meng & Takeichi, 2009; Ebrahim et al., 2013). In non-muscle cells, including epithelial cells, it is assumed that intracellular Ca^2+^ ion (iCa^2+^)-dependent phosphorylation of Myosin Regulatory Light Chain (MRLCs) generates mechanical force as seen in smooth muscle cells (Turner et al., 2024; Hathaway & Adelstein, 1979; Garrido-Casado et al., 2024). The stimulation of contractile force is released by opposing MRLC phosphatases (Watanabe et al., 2007; Bresnick, 1999). Both myosin and its upstream regulators, Rho-kinase Coiled-coil Kinase (ROCK), Myosin Light Chain Kinase (MLCK), and Myosin Light Chain Phosphatase (MLCP) are required for proper actomyosin contraction dynamics (Totsukawa et al., 2000; Kassianidou et al., 2017). Changes to the actomyosin contractile network have been shown to change epithelial cell morphology; in particular, this network can be used to drive apical constriction and tissue morphogenesis (Ranie & White, 2025; Martin et al., 2009; Chugh & Paluch, 2018; Martin & Goldstein, 2014; Lemke et al., 2021; Clarke & Martin, 2021). It is also unknown whether iCa^2+^-dependent phosphorylation of MRLCs is required for regulation of a flat apical surface of a steady-state epithelium.

Using Madin-Darby Canine Kidney (MDCK) cell monolayers, the present study experimentally probed the mechanisms underlying the flatness of the apical epithelial surface, distinctly from the overall epithelial flatness. Here we show the flatness of an epithelial sheet was preserved after growth on irregular non-uniform basement matrices. We also reveal that pharmacological and molecular perturbations disrupting myosin activation caused a loss of apical surface flatness that was more severe than perturbations of the actin networks. Finally, we show that epithelial cells transiently, and asynchronously, evoked iCa^2+^ fluxes across the sheet, implying that calcium is available for a potential activation of MLCK-mediated mechanical force. Our data reveal that apical and basal cellular surfaces are not tightly coordinated, but independently regulated, and hint that myosin may have a role in the apical morphology of epithelial cells.

## Results

### Definition of flatness using confluent MDCK monolayers

To evaluate epithelial architecture, we used MDCK cells, an established model for epithelial sheet formation (Dukes et al., 2011; Wells et al., 2013; Misfeldt et al., 1976), and immunofluorescence labeling of polarity markers. Individual cell and apical height was quantified at both the tissue-level (macro-height) and cellular-level (micro-height) for MDCK confluent monolayers to define flatness. To assess the loss of flatness, three parameters were considered: 1) height differences between immediately adjacent cells, 2) the overall distribution of cell heights within a defined area, and 3) apical domain height of individual cells. These metrics were applied for macro- and micro-height measurements to characterize surface irregularity, or loss of apical flatness. In cases where the distributional analysis (parameter 2), and apical domain height (parameter 3) sufficiently capture variation in height and spatial heterogeneity, the pairwise height difference data (parameter 1) were omitted without compromising the evaluation of flatness.

Immunofluorescence for β-catenin and ZO-1 was used to define lateral and apical junctions, respectively, and phalloidin to define cell boundaries for measuring macro- and micro-height (Figure 1 A, A’). Our confocal 3D reconstruction analysis revealed that confluent MDCK cells showed stereotypical accumulation patterns of ZO-1, β-catenin, and phalloidin-bound actin bundles (Figure 1 B), consistent with previous studies (McNeil et al., 2006; Kuo et al., 2022; Leng et al., 2020; Odenwald et al., 2018; Fanning et al., 2012).

**Figure 1.**
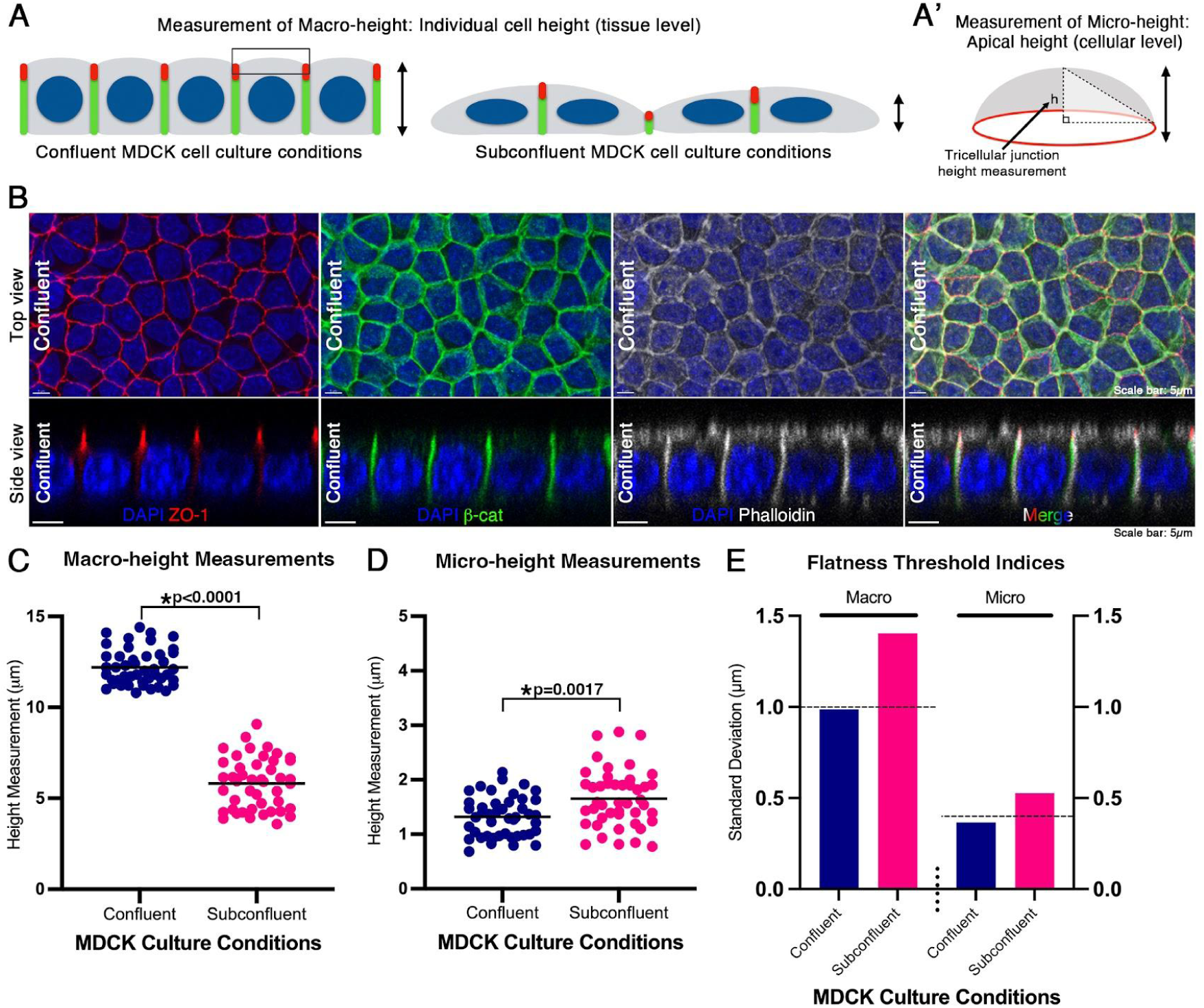
Definition of flatness using confluent MDCK monolayers. (A) Schematic showing the measurements of macro-height of individual MDCK cell height when grown in confluent and subconfluent cell culture conditions. (A’) Micro-height or apical cell height was measured from the apex of filamentous actin staining to the base of the tricellular junctions. (B) Top and side views of stacked confocal optical sections of a MDCK confluent monolayer immunostained with ZO-1 (red), β-catenin (green), with filamentous actin dye, phalloidin (grey), and a DNA counterstain, DAPI (blue). (C) Average cell height or macro-height measurements for confluent MDCK cells was 12.21 µm ± 0.99 µm s.d., and 5.81 µm ± 1.40 µm s.d. for subconfluent cells (U=0.00, n_1_=45, n_2_=45, p<0.0001). (D) Same as (C) but of apical height or micro-height: 1.32 µm ± 0.37 µm s.d. for confluent cells, and 1.65 µm ± 0.53 µm s.d. for subconfluent cells (U=628, n_1_=45, n_2_=45, p=0.0017). Conditions were statistically compared with a non-parametric Mann-Whitney U test. Unequal variance was confirmed with Levene’s test (p=0.0093 for macro-height, p=0.0274 for micro-height). (E) Standard deviations of macro- and micro-height of confluent and subconfluent monolayers. Flatness threshold indices are indicated with horizontal dashed lines. (Scale bars: 5µm)

Trans-epithelial electrical resistance (TEER) measurements of MDCK cells were used to determine when confluent monolayers formed (Srinivasan et al., 2015; Schimetz et al., 2024). MDCK monolayers displayed the highest electrical resistance of 279.6 Ω*cm^2^ ± 33.23 Ω*cm^2^ s.d. at 45 hours over a three-day measurement period of the same monolayer (sFigure 1 A), similar to other reported TEER measurements for MDCK cells (Fedi et al., 2021). The data demonstrate that in our hands, 45 hours was necessary to define a confluent monolayer on which we could complete our analysis (sFigure 1 A). Macro-height data was collected as a linear measure of the cell height from basement membrane to the ZO-1 junctions (Figure 1 A, sFigure 1 B, C). Micro-height was determined by measuring a degree of height difference between the midpoint of the phalloidin-positive apical surface, and ZO-1-positive tricellular junctions (Figure 1A’, sFigure 1 B, C). The average macro-height and micro-height measurements for the confluent MDCK monolayer were 12.21 μm ± 0.99 μm standard deviation (s.d.), and 1.32 μm ± 0.37 μm s.d., respectively (n=45 cells) (Figure 1 C, D). The standard deviation was used to generate a macro-height and micro-height index, and served as a threshold to evaluate if overall flatness was lost (Figure 1 E).

To validate the threshold indices, we used subconfluent MDCK cells to confirm the expected morphology and reflect a loss of flatness. Subconfluent MDCK cells remain scattered and do not yet form a cohesive monolayer, providing a control for testing the validity of the threshold indices (sFigure 1 D). Subconfluent MDCK cells displayed a higher degree of variation in both macro- and micro-height as compared to confluent monolayers (Figure 1 B-E, and sFigure 1D). For example, the s.d. of macro-height for subconfluent cells was 1.40 μm and exceeded the threshold of a confluent monolayer, 0.99 μm s.d. (Figure 1 C, E). The s.d. of micro-height for subconfluent cells was 0.53 μm s.d., and also exceeded the threshold of confluent monolayers, 0.37 μm s.d. (n=45 cells) (Figure 1 D, E). Confluent monolayers and subconfluent cells were statistically compared with a Mann-Whitney U-test (p<0.0001 for macro-height and p=0.0017 for micro-height), and unequal variance was confirmed with Levene’s test (p=0.0093 for macro-height and p=0.0274 for micro-height). Based on these results, the threshold indices using confluent monolayers were confirmed to define a loss of flatness (Figure 1 E). MDCK cell samples with s.d. ≥ 1.0 µm for macro-height, and s.d. ≥ 0.4 µm for micro-height were considered to lose flatness (Figure 1 E).

In summary, these data show that confluent MDCK monolayers display a flat, uniform morphology that can be described using s.d. threshold indices. These threshold indices can be used to define a loss of flatness. In the following experiments, we examined the biophysical and molecular processes underlying the maintenance of monolayer flatness.

### Association between apical flatness and the ECM

Previous studies have revealed that lateral factors within cells as well as stiffness of the basal surface can function to regulate cell height (Kondo & Hayashi, 2015; Cai & Mostov, 2012; Töpfer, 2023; Perez-Tirado et al., 2025; Janmey et al., 2020; Tang, 2017). The ECM regulates many aspects and properties of epithelial sheet morphogenesis such as apico-basal polarity, cell proliferation, differentiation, and death; serving to establish and maintain proper tissue function (Perez-Tirado et al., 2025; Diaz-de-la-Loza et al., 2018; Kozyrina et al., 2020; Frantz et al., 2010; Rozario & DeSimone, 2010; Muncie & Weaver, 2018). However, it is unknown whether the ECM plays a role in defining the apical surface geometry of steady-state epithelial sheets.

To test this possibility, we engineered a basement membrane with a patched deposition of fluorescently labeled laminin on transwell filters to represent an uneven ECM (Figure 2 A). Under our culture conditions, the patched laminin still allowed MDCK cells to successfully establish a confluent monolayer (n=6 samples). Laminin patches were classified as ‘steep’ or ‘shallow’ based on the ratio of apical to basal surface area. Steep laminin patches had a ratio of 2.35 ± 0.26 s.d for apical/basal surface area. Shallow laminin patches had a ratio of 1.74 ± 0.31 s.d. for apical/basal surface area (Figure 2 B). Both types of epithelium were clearly delineated by a laminin patch ratio that was either larger (steep) or smaller (shallow) than 2 (n=16 laminin patches) (Figure 2 B).

**Figure 2.**
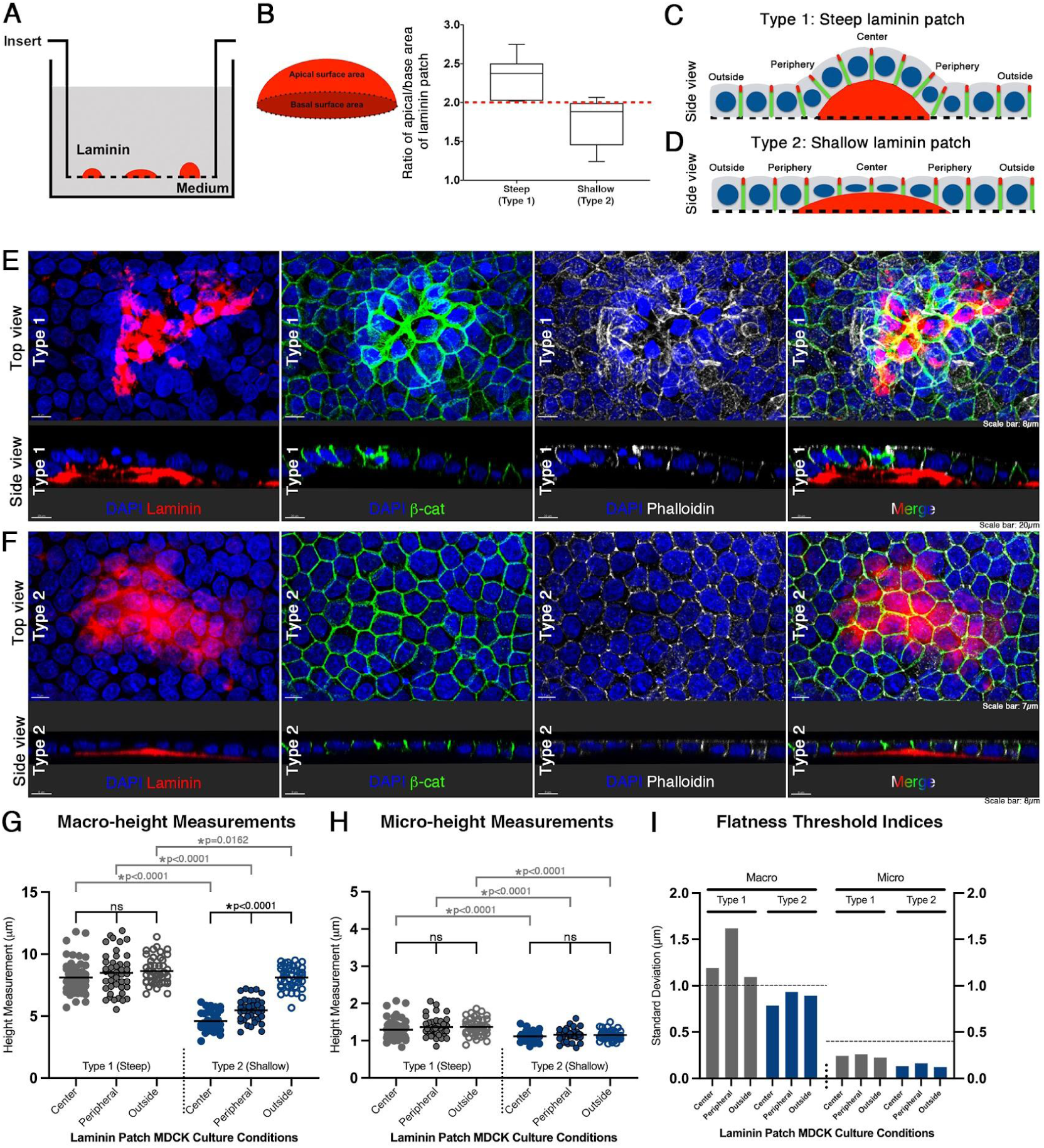
Association between apical flatness and the extracellular matrix (ECM). (A) Schematic of Rhodamine-laminin patches on Transwell membranes prior to MDCK cell culture. (B) Quantification of steep and shallow Rhodamine-laminin patches. The ratio of apical/basal area of the laminin patch delineated the two categories of steep and shallow patches. ‘Steep’ patches had a ratio of 2.35 ± 0.26 s.d., and ‘shallow’ patches had a ratio of 1.74 ± 0.31 s.d. Note: Both steep/shallow patches were clearly delineated by a ratio that was either > or < 2, with the ‘steep’ patch ratio > 2, and ‘shallow’ patch ratio < 2 (n=16 laminin patches). (C, C’) Two types of epithelial response in morphology based on the steepness of the laminin patch. Cells with complete, partial, or little to no basement membrane positioned on the patch were defined as center, peripheral, or outside, respectively. (D) Top and side views of stacked confocal optical sections of a Type 1 MDCK confluent monolayer immunostained with β-catenin (green), and filamentous actin dye, phallodin (grey), and a DNA DAPI (blue) counterstain when grown on a ‘steep’ Rhodamine-laminin patch (red). (E) As in (D) but of a Type 2 MDCK confluent monolayer growing on a ‘shallow’ Rhodamine-laminin patch (red). (F) Macro-height measurements for Type 1 (grey) and Type 2 (blue) monolayers. Individual cell height analysis was performed based on the relative position on the laminin patch. Average macro-height of center, peripheral, or outside cells was 8.10µm, 8.42 µm, and 8.64, respectively (n=149 cells). For Type 2 cells, average macro-height for center, peripheral, and outside cells was 4.60 µm s.d., 5.47 µm, and 8.12 µm, respectively (n=117 cells). (G) As in (F), but of micro-height measurements. For Type 1 cells, average micro-height measurements for center, peripheral, and outside cells were 1.29µm, 1.36 µm, and 1.37 µm, respectively (n=149 cells). For Type 2 cells, average micro-height for the center, peripheral, and outside cells was 1.12µm, 1.16 µm,, and 1.15 µm, respectively (n=117 cells). (H) Standard deviations of macro- and micro-height of Types 1 (steep) and 2 (shallow) MDCK confluent monolayers. Flatness threshold indices established in Figure 1 are indicated with horizontal dashed lines. Note: Type 1 epithelium loses flatness, while Type 2 epithelium maintains flatness by modulating individual cellular behaviors on the laminin patch. (Scale bars: 8µm, 20µm, 7µm)

Epithelia were classified into Type 1 or 2. Type 1 epithelium was associated with steep laminin patches, while Type 2 was associated with shallow laminin patches (Figure 2 B-D). Type 1 epithelium displayed a wavy surface tissue morphology that traveled along the contours of the steep laminin patch (n=3 samples) (Figure 2 C, E). Type 2 epithelium had a flat tissue geometry that modulated to adapt along the shallow laminin patches (n=3 samples) (Figure 2 D, F).

To determine the flatness of the two types of epithelia, individual cell height analysis was performed based on the relative position on the laminin patch. Cells with complete, partial, or little to no basement membrane positioned on the patch were defined as center, peripheral, or outside, respectively (Figure 2 C, D). We quantified macro- and micro-height using β-catenin staining (sFigure 1).

Macro-height measurements of Type 1 (steep) epithelial cells of center, peripheral, or outside were 8.10µm, 8.42 µm, and 8.64 µm, respectively (n=149 cells) (Figure 2 G). A Kruskal-Wallis statistical test was performed to compare mean macro-height measurements between groups of Type 1 cells (H(2, 153)=5.097, p=0.0782), with unequal variance confirmed using Levene’s statistical test (p=0.0177). Average micro-height measurements for center, peripheral, and outside cells were 1.29µm, 1.36 µm, and 1.37 µm, respectively (Figure 2 H). A One-way ANOVA statistical test was performed to compare mean micro-height measurements between groups of Type 1 cells (F (2, 146)=1.537, p=0.2185), with equal variance confirmed using Levene’s statistical test (p=0.8414). Average macro- and micro-height measurements were not statistically different between center, peripheral, and outside cells of Type 1 epithelium (Figure 2 G, H).

Although average macro-heights for Type 1 cells were similar (Figure 2 G), the s.d values are above the threshold index of flatness of 1.0µm, indicating that flatness was lost (Figure 2 I). These data demonstrate that Type 1 epithelium loses its flatness by developing increased cell height variability among central, peripheral, and outside cells.

For Type 2 (shallow) epithelium, average macro-height for center, peripheral, and outside cells was 4.60 µm, 5.47 µm, and 8.12 µm, respectively (n=117 cells) (Figure 2 G). The data show that macro-height is significantly different between center, peripheral, and outside cells in Type 2 epithelia (One-way ANOVA, F(2, 114)=177.5, p<0.0001; Levene’s test, p=0.7065). However, the average micro-height measurements were not significantly different for the center, peripheral, and outside cells was 1.12µm, 1.16 µm, and 1.15 µm, respectively (One-way ANOVA, F(2, 114)=0.9336, p=0.3961; Levene’s test, p=0.5907) (Figure 2 H). The s.d. of macro-and micro-height of Type 2 cells did not exceed the flatness threshold indices, indicating that the monolayer remained flat (Fig. 2 I). These results reveal a striking pattern: Type 2 epithelia on a shallow laminin patch adapt to maintain a uniform tissue flatness by decreasing their individual cell height, or macro-height.

In summary, these data show that flatness is lost in Type 1 epithelium on a steep laminin patch, and is maintained for Type 2 epithelium on a shallow laminin patch. Measurements of macro- and micro-height for Type 2 center, peripheral, and outside cells were significantly shorter than Type 1 counterparts confirming the two distinct monolayer behaviors defined by steep/shallow laminin patches (six comparisons, four Mann-Whitney U tests (U=563, 378, 407.5, and 60, all p<0.0001) and two Student’s t-tests (t=15.54, p<0.0001; t=2.452, p=0.0162) (Figure 2 G, H). The apical surface of the two different types of epithelial sheet was continuous regardless of basement membrane evenness. These observations suggest that it is not the ECM component, but rather the steepness and topography of the ECM that determines whether the steady-state epithelial sheet maintains a constant height along the patch, or decreases to adapt to laminin topography. These data support that apical surface flatness is differentially regulated by the basement membrane after a steepness threshold. Our findings suggest that individual MDCK cells can detect substrate steepness, and respond by regulating their height at both macro- and micro-levels.

To identify potential intracellular factors underlying these differential adaptive responses in MDCK cells, we next examined whether cytoskeletal components contribute to the regulation of cell height.

### Pharmacological inhibition to screen cytoskeleton fibers for potential involvement in macro- or apical micro-height

To test whether cytoskeletal elements contribute to the regulation of epithelial monolayer flatness, MDCK monolayers were treated with pharmacological inhibitors targeting actin (cytochalasin D), intermediate filaments (withaferin A), and microtubules (nocodazole) (Figure 3 A, B). Treated cells were then fixed and labeled with ZO-1, β-catenin, and phalloidin to visualize epithelial architecture (Figure 3 C-G). Control cells incubated with DMSO showed indistinguishable patterns of polarity markers from untreated MDCK monolayers (n=45 cells each) (Figure 3 C, D). This was also consistent with macro- and micro-height mean measurements using the Kruskal-Wallis/Dunn’s test and s.d. of macro- and micro-height was below the threshold indices (H(5, 210)=82.31 for macro, 57.48 for micro) (Figure 3H, I).

**Figure 3.**
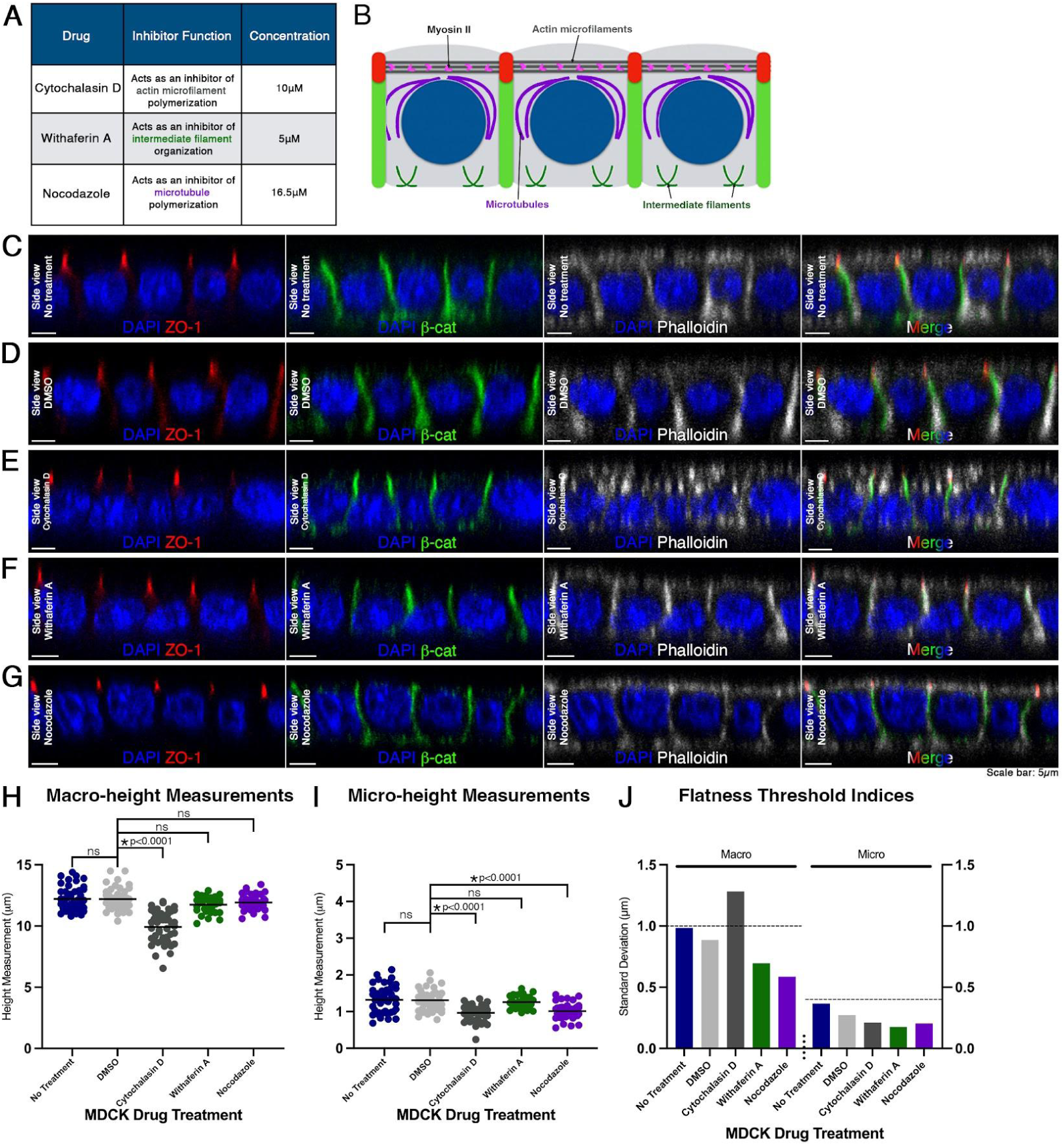
Pharmacological inhibition to screen cytoskeleton fibers for potential involvement in macro- or apical micro-height. (A) Table showing the function and concentration of drugs used to inhibit cytoskeletal polymerization; cytochalasin D, withaferin A, and nocodazole. (B) Schematic of MDCK cells showing localization of actin microfilaments, intermediate filaments, and microtubules. (C) A side view of untreated MDCK cells immunostained with ZO-1 (red), β-catenin (green), and filamentous actin dye, phalloidin (grey), and DNA DAPI (blue) counterstain. (D) As in (C) but cells treated with DMSO, (E) cytochalasin D, (F) withaferin A, and (G) nocodazole. (H) Measurements of macro-height for individual MDCK cells (n=45 cells for each drug treatment). Average macro-height for cells with No Treatment, or treated with DMSO, cytochalasin D, withaferin A, and nocodazole were 12.21 µm ± 0.99 µm s.d., 12.20 µm ± 0.89 µm s.d., 9.93µm ± 1.28 µm s.d., 11.75 µm ± 0.70 µm s.d., and 11.92 µm ± 0.59 µm s.d., respectively. (I) As in (H) but of micro-height. Average micro-height for No Treatment, DMSO, cytochalasin D, withaferin A, and nocodazole was 1.31 µm ± 0.27 µm s.d., 1.32 µm ± 0.37 µm s.d., 0.97µm ± 0.21 µm s.d., 1.26 µm ± 0.18 µm s.d., and 1.01 µm ± 0.20 µm s.d., respectively. (J) Flatness threshold indices of macro- and micro-height of treated and control MDCK confluent monolayers. Note: Only cytochalasin D treatment of the MDCK monolayer resulted in a loss of flatness. (Scale bars: 5µm)

In cytochalasin D-treated cells, ZO-1 and β-catenin localization remained indistinguishable from DMSO control groups (Figure 3E). However, phalloidin staining revealed disruption of filamentous actin into small aggregates, as compared to the uniform and stereotypical appearance in DMSO controls (Figure 3E) (Stevenson and Begg, 1994; Schliwa, 1982). Cytochalasin D-treated cells also displayed disruption of apical and basal actin, and reduced macro- and micro-height (p<0.0001) (Figure 3 E, H, I). A decrease in cell height has also been previously reported (Stevenson & Begg, 1994; Mortensen & Larsson, 2003; Wells et al., 1998). The s.d. of macro-height exceeded the threshold index indicating that treatment with cytochalasin D results in loss of monolayer flatness (Figure 3 J).

Withaferin A-treated and nocodazole-treated cells exhibited no apparent changes in fluorescent staining localization, with ZO-1, β-catenin, and phalloidin staining indistinguishable from DMSO controls (Figure 3F, G). Neither of these treatments created significant changes in macro-height (Figure 3 H); however, micro-height of nocodazole-treated cells was decreased (Figure 3 I) (p<0.0001). The s.d. for macro- and micro-height for cells treated with withaferin A or nocodazole were below the threshold indices (Figure 3 J). This result shows that inhibiting microtubule polymerization with nocodazole may disrupt micro-height, without affecting the overall flatness of the epithelial sheet.

These data support the idea that actin plays a role in regulating cell height, or macro-height, while both actin and microtubules contribute to apical micro-height. Actin filaments and microtubules are both indispensable for the function of their partner motor proteins, myosin and dynein, respectively, together enabling many cellular processes including cell shape regulation (Huang et al., 2022; Quintin et al., 2008; Pollard & Cooper, 2009; Oelz et al., 2018; Barlan & Gelfand, 2017). In our hands, no recognizable role in MDCK monolayer architecture was identified for intermediate filaments.

### Defective flatness by inhibition of myosin II

We next examined the role of myosin II, the canonical actin partner in the cellular contractile network, in the maintenance of steady-state epithelial flatness. We perturbed the actomyosin contractile network, applying blebbistatin (myosin II ATPase), Y27632 (ROCK), and ML-7 (MLCK) to MDCK monolayers to compromise myosin II motor protein complex function (Watanabe et al., 2007; Kovács et al., 2004; Goeckeler et al., 2008; Vasquez et al., 2014; Cui et al., 2010) (Figure 4 A, B).

**Figure 4.**
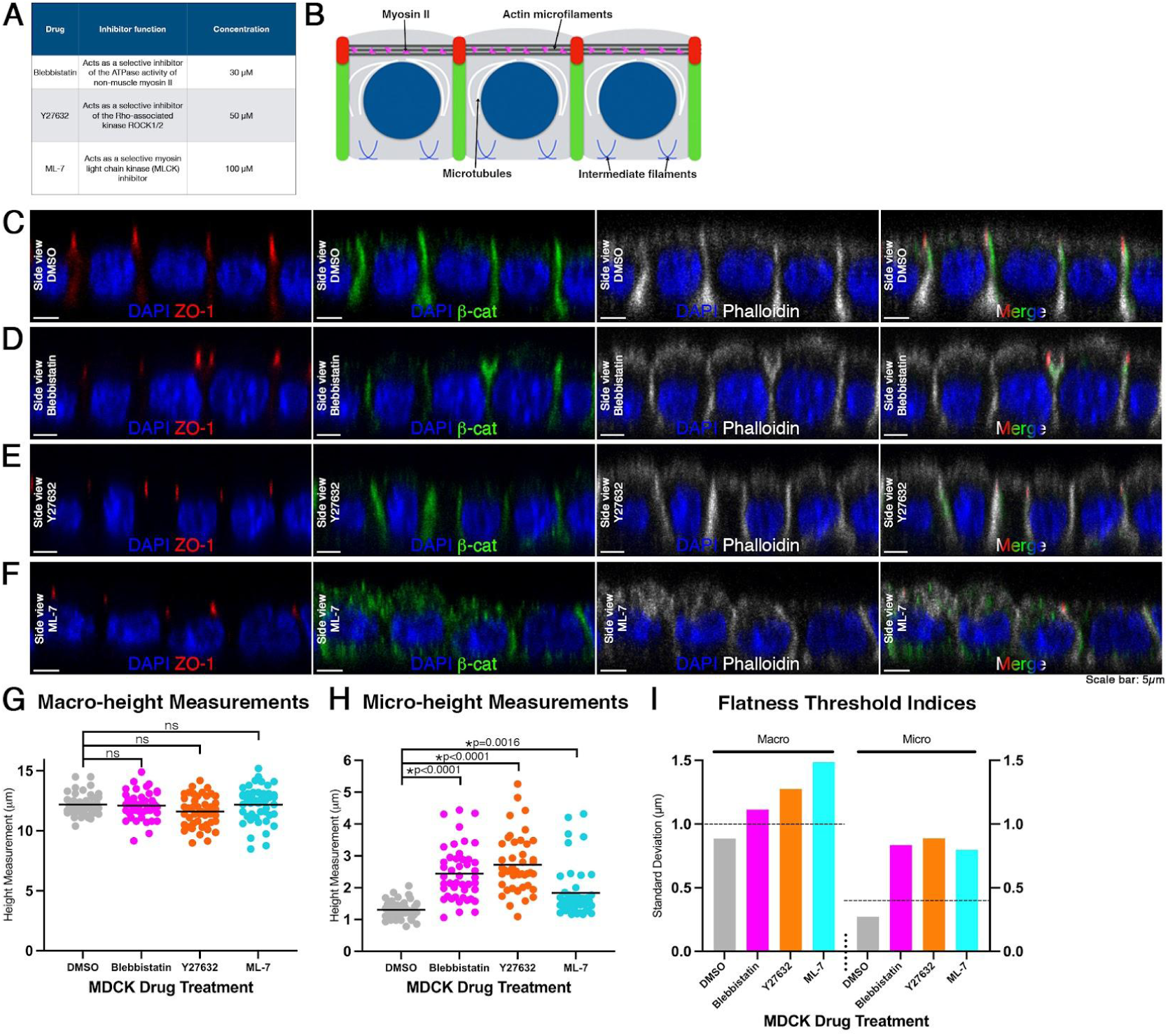
Defective flatness by inhibition of myosin II. (A) Table showing drugs used to target myosin II, inhibitor function, and concentration. (B) Schematic showing the localization of myosin II interaction with actin in an MDCK monolayer. (C) A side view of a single optical section of a MDCK confluent monolayer treated with DMSO immunostained with ZO-1 (red), β-catenin (green), and filamentous actin dye, phalloidin (grey), and DNA DAPI (blue) counterstain. (D) As in (C) but treated with 30 µM blebbistatin, 50 µM Y27632 (E), and 100 µM ML-7 (F). (G) Individual cell measurements of macro-height for DMSO and drug-treated MDCK monolayers. Average macro-height for DMSO, blebbistatin, Y27632, and ML-7-treated cells was 12.20 µm ± 0.89 µm s.d., 12.11 µm ± 1.11 µm s.d., 11.61 µm ± 1.28 µm s.d., 12.20 µm ± 1.49 µm s.d., respectively (n=45 cells for each treatment). (H) As in (G) but of micro-height. Average micro-height for DMSO, blebbistatin, Y27632, and ML-7-treated cells was 1.30 µm ± 0.27 µm s.d., 2.44 µm ± 0.84 µm s.d., 2.72 µm ± 0.89 µm s.d., 1.84 µm ± 0.80 µm s.d., respectively. (I) Flatness threshold indices of macro-and micro-height of treated and control MDCK confluent monolayers. Note: All myosin II inhibitor treatments resulted in a loss of flatness. (Scale bars: 5µm)

After treatment with blebbistatin or Y27632, ZO-1 and β-catenin localization patterns remained indistinguishable from DMSO controls (Figure 4 C-E). Treatment with ML-7 resulted in β-catenin localization throughout the cell rather than at cell junctions, differing from DMSO controls (Figure 4 F). Individual cells for all three treatments showed an increased apical domain, and a dome-like spherical cap of actin as compared to flat apical actin as in control DMSO samples (Figure 4 C-F). Average macro-height of cells treated with blebbistatin, Y27632, or ML-7 showed no statistical difference from controls (Figure 4 G). However, blebbistatin or Y27632-treated cells exhibited significantly greater average micro-height values compared to DMSO controls (H(4, 180)=87.19, p<0.0001; Dunn’s post-hoc test, p<0.0001 for blebbistatin and Y27632) (Figure 4 H). ML-7-treated cells showed a slightly milder phenotype, yet average micro-height still differed significantly from DMSO (Dunn’s post-hoc test, p=0.0016) (Figure 4 H). For all three treatments, the flatness threshold indices were exceeded for both macro- and micro-height (Figure 4 I). These data show that inhibiting myosin II ATPase with blebbistatin, or ROCK function with Y27632, or MLCK function with ML-7 results in overall loss of MDCK monolayer flatness.

The observation of apical domes with our myosin drug perturbations indicate that some of the underlying actin cytoarchitecture is disorganized. An important component of the apical morphology of MDCK cells is microvilli (Klingner et al., 2014; Sharkova et al., 2023; Gaeta et al., 2021). Microvilli are actin filament-based structures that lack myosin II along their length, with myosin II instead enriched in the underlying filamentous network, or terminal web (Mooseker & Tilney, 1975; Chinowski et al., 2020; Hirokawa et al, 1982). To investigate whether microvilli are disorganized in our MDCK monolayers treated with ML-7, the myosin II treatment that resulted in the mildest phenotype of apical domes, we performed Transmission Electron Microscopy (TEM). TEM analysis revealed a decrease in microvilli number for ML-7-treated MDCK cells as compared to DMSO controls (sFigure 2 A, B). ML-7-treated cells had an average of 6.75 ± 3.54 s.d. microvilli, while DMSO controls had an average of 35.88 ± 6.13 s.d. microvilli (n=8 cells for each treatment) (t=11.64, p<0.0001) (sFigure 2 C). As actin-based microvilli decreased in ML-7-treated cells, these results suggest a potential non-contractile role for myosin II, where perturbations of MLCK activity result in loss of flatness via microvilli (Lombardo et al., 2022; Chinowski et al., 2020).

We recognize that pharmacological treatments may induce off-target effects, and thus the observed outcomes should be interpreted carefully, as they may not be exclusively attributable to the intended actions of the drugs. Therefore, precise genetic perturbations of myosin II are necessary.

### Transgenic perturbation of myosin II alters flatness

To investigate myosin II roles in the actomyosin contractile complex to regulate flatness, we caused an imbalance in contractile dynamics. MLCP has both a regulatory/myosin targeting subunit (MYPT1) and a catalytic subunit (PP1δ) (Koga & Ikebe, 2008; Shibata et al., 2014). The *MYPT1^T696A;T853A^* mutant has two point mutations, T696A and T853A, in the catalytic region of the PP1δ, therefore MLCP is locked in its active conformation (Koga & Ikebe, 2008; Álvarez-Santos et al., 2020). Constitutively active MLCP will cause MLC inactivation, and result in an imbalance of relaxation and contraction of myosin (Khasnis et al., 2014; Khromov et al., 2009). We used a transient transfection model for introducing the *MYPT1^T696A;T853A^* variant into MDCK cells and comparing mutant and wild-type cells.

Transiently transfected MYPT1^wt^-GFP MDCK cells exhibited flatness within the monolayer and were used as controls (Figure 5 A). Control MYPT1^wt^-GFP cells were categorized as GFP(+) cells that expressed the plasmid and GFP, and GFP(-) cells that did not express the plasmid. Macro- and micro-height measurements showed no significant difference between GFP(-) and GFP(+) cells, and s.d. threshold indices were not exceeded (Figure 5 C-E). These data show that MDCK cells transiently transfected with our control MYPT1^wt^-GFP plasmid retained flatness, and served as an appropriate control for comparison of the mutant *MYPT1^T696A;T853A^* cell line.

**Figure 5.**
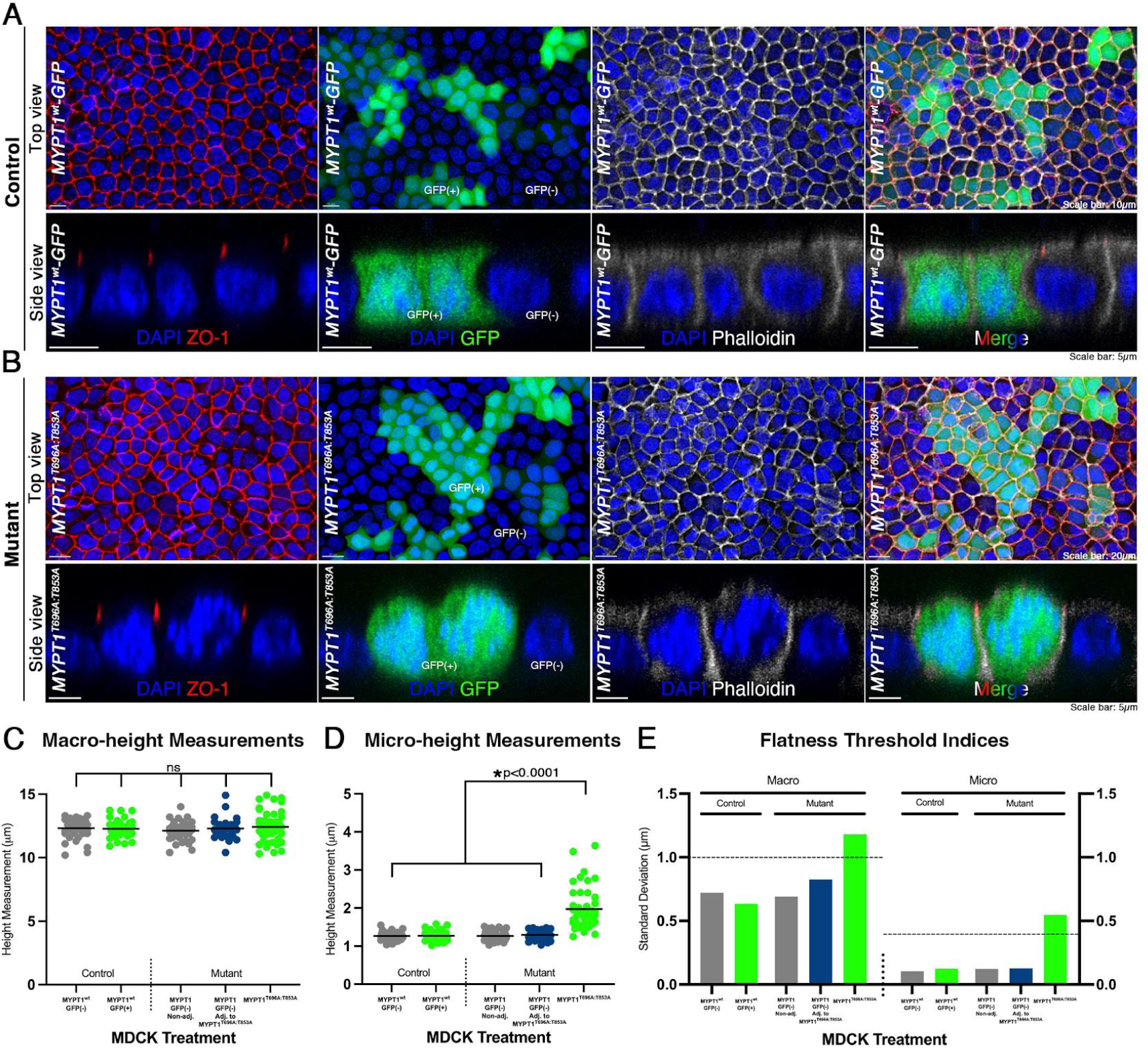
Transgenic perturbation of myosin II alters flatness. (A) Top and side views of stacked confocal optical sections of a control MDCK confluent monolayer transfected with MYPT1^wt^-GFP (green) immunostained with ZO-1 (red), and filamentous actin dye, phalloidin (grey), and DNA DAPI (blue) counterstain. (B) As in (A) but of *MYPT1^T696A;T853A^* mutant MDCK lines. (C) Individual measurements of macro-height for MYPT1^wt^-GFP control cells (left of vertical dashed line), and *MYPT1^T696A;T853A^*mutant cells (right of vertical dashed line). Average macro-height measurements for MYPT1^wt^-GFP control cells was 12.30 µm ± 0.77 µm s.d., 12.46 µm ± 0.71 µm s.d., and 12.27 µm ± 0.63 µm s.d. for GFP(-), GFP(-) adjacent to GFP(+), and GFP(+) cells, respectively. Average macro-height for *MYPT1^T696A;T853A^* cells was 12.20 µm ± 0.65 µm s.d., 12.41 µm ± 0.80 µm s.d., and 12.42 µm ± 1.18 µm s.d. for GFP(-), GFP(-) adjacent to *MYPT1^T696A;T853A^* and GFP(+) *MYPT1^T696A;T853A^* respectively. (D) As in (C) but of micro-height for control MYPT1^wt^-GFP cells 1.27 µm ± 0.10 µm s.d., 1.27 µm ± 0.08 µm s.d., and 1.27 µm ± 0.12 µm s.d., respectively. Average micro-height for mutant *MYPT1^T696A;T853A^* cells was 1.34 µm ± 0.15 µm s.d., 1.38 µm ± 0.14 µm s.d., and 1.97 µm ± 0.55 µm s.d., respectively. A Kruskal-Wallis statistical test was performed to compare mean macro- or micro-height between all groups (H(5, 210)=93.94, p=0.4555 for macro-height, p<0.0001 for micro-height; Dunn’s post-hoc test, p<0.0001 for all treatments compared to *MYPT1^T696A;T853A^*). Unequal variance was confirmed using Levene’s test (p=0.0010 for macro-height, p<0.0001 for micro-height). (E) Flatness threshold indices of macro- and micro-height of treated and control MDCK confluent monolayers. Note: Only mutant *MYPT1^T696A;T853A^* cells resulted in a loss of flatness. GFP(-) cells adjacent to *MYPT1^T696A;T853A^* cells remained flat, suggesting that maintaining flatness is a cell autonomous event. (Scale bars: 10µm, 20µm, 5µm)

We divided the monolayers treated with the mutant plasmid into three groups: (1) GFP(+) *MYPT1^T696A;T853A^* cells expressing the plasmid with both point mutations, (2) adjacent GFP(-) cells not expressing the plasmid, and (3) non-adjacent GFP(-) cells also lacking plasmid expression. Similar to cells treated with myosin II complex inhibitors (Figure 4), mutant *MYPT1^T696A;T853A^*cells showed an increased apical domain and dome-like spherical actin caps (Figure 5 B); while both adjacent and non-adjacent GFP(-) cells showed a similar flat morphology to our controls (Figure 5 A, B). Average macro-height of *MYPT1^T696A;T853A^* cells showed no statistical difference from adjacent or non-adjacent GFP(-) cells, nor from GFP(+) and GFP(-) MYPT1^wt^-GFP control cells (Figure 5 C). *MYPT1^T696A;T853A^* cells exhibited significantly increased micro-height compared to adjacent and non-adjacent GFP(-) cells, as well as control GFP(+) and GFP(-) cells (H(5, 210)=93.94, p<0.0001; Dunn’s post-hoc test, p<0.0001 for all treatments compared to *MYPT1^T696A;T853A^*) (Figure 5 D). Loss of flatness and an increased apical domain was observed exclusively in *MYPT1^T696A;T853A^*cells, whereas all other groups remained flat (Figure 5 A, B, E). Taken together, the loss of flatness for genetic and drug disruptions of MLCP and MLCK (Figure 4), respectively, provide supportive evidence that myosin II morphodynamics may regulate flatness in a steady-state epithelium.

These data also show that alterations to macro- or micro-height by genetic disruption of MLCP dynamics are limited to the transfected cells, and do not significantly influence adjacent cells. Transient transfection creates a mosaic of plasmid-expressing and non-expressing cells within the same monolayer, enabling us to assess whether cellular perturbations act autonomously or influence neighboring cells, non-autonomously (Figure 5). The results indicate that forces maintaining MDCK cell flatness are cell-autonomous events, revealing that individual cells may independently govern their own architecture. Our findings suggest that individual MDCK cells may autonomously regulate flatness, providing insights into the underlying mechanisms that contribute to the maintenance of flatness at the tissue- and cellular-level.

### Intracellular Ca^2+^ (iCa^2+^) and maintenance of apical flatness in a steady-state epithelium

Intracellular calcium (iCa^2+^) is an important regulator of many signaling pathways and cellular functions (Bagur & Hajnóczky, 2017; Moccia et al., 2023; Bers, 2007). In particular, iCa^2+^ is a key secondary messenger that helps govern cytoskeletal function, including the actomyosin network, in a number of tissues and organ systems (Kuo & Ehrlich, 2015; Yerna et al., 1979; Christodoulou & Skourides, 2015; Antunes et al., 2013; Suzuki et al., 2017; Castillo et al., 1998; Sahu et al., 2017). It is unknown whether iCa^2+^ abundance in an epithelial cell contributes to the maintenance of apical flatness.

Therefore, we examined changes in iCa^2+^ in individual cells in our MDCK monolayers. We took advantage of a calcium indicator dye, Fura-2, to perform fluorometric calcium imaging (Barreto-Chang & Dolmetsch, 2009). The fluorescence intensity of the ratio of Fura-2 340/380nm can be used to calculate iCa^2+^ concentrations (Paredes et al., 2008). Real-time calcium imaging showed rapid, intermittent calcium transients with high [iCa^2+^] appearing as red fluorescence and low [iCa^2+^] as green (Figure 6 A, sMovie 1). With [iCa^2+^] tracked over time, the imaging revealed asynchronous calcium oscillations among adjacent cells in the monolayer (n=6 cells, 3 samples total) (Figure 6 A-C, sMovie 1). The asynchronous iCa^2+^ oscillations across the monolayer show that individual cells regulate their [iCa^2+^] in an uncoordinated and independent manner from juxtaposed cells. These iCa^2+^ oscillations may work together with cell autonomous height regulation via the myosin II complex, creating a model where increased [iCa^2+^] stimulates myosin contraction nonsynchronously among adjacent cells (Figure 6 D). The data support a model of spatio-temporal asynchronous iCa^2+^ oscillations that may play a role in balancing actomyosin dynamics across the monolayer to maintain flatness.

**Figure 6.**
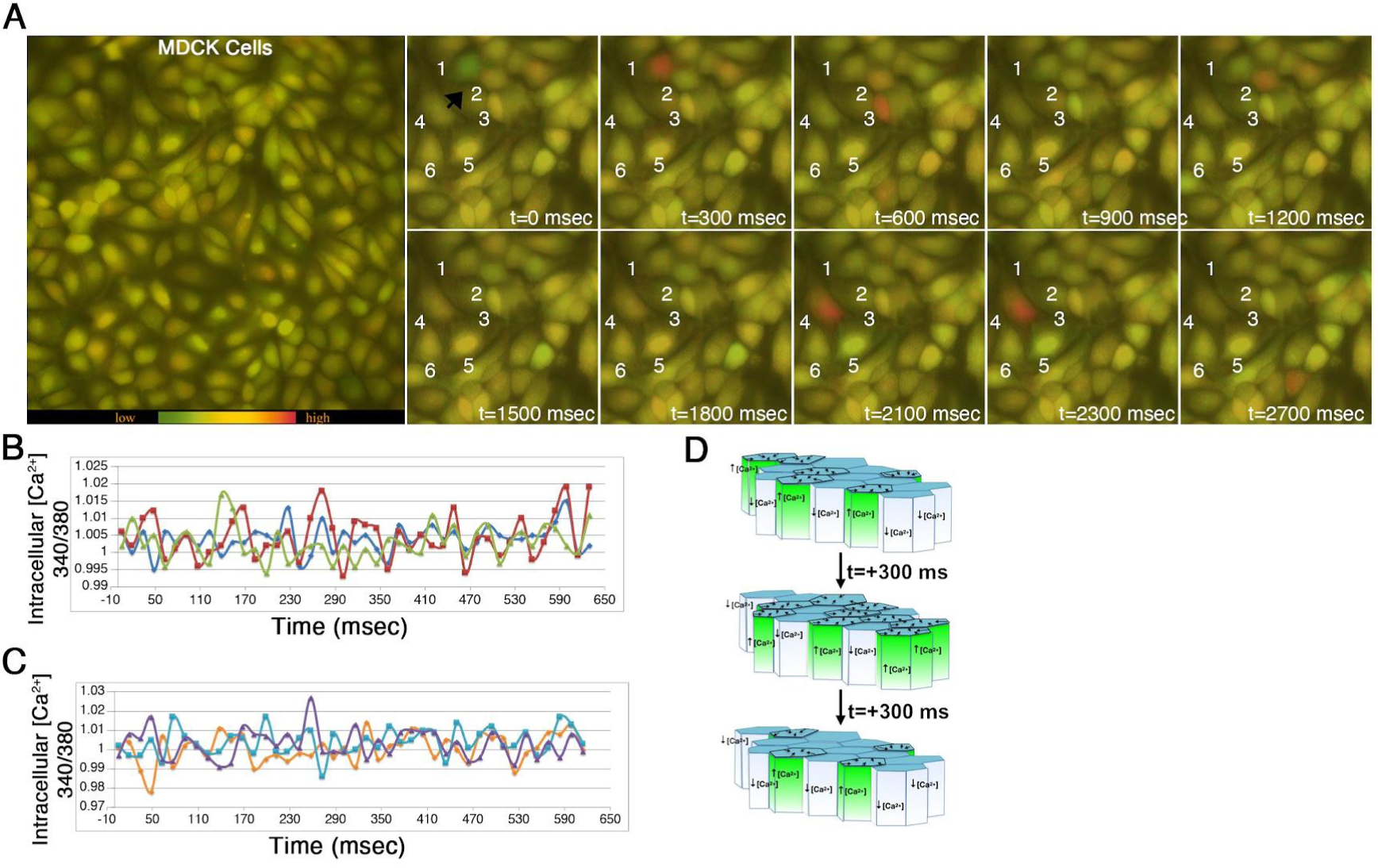
Intracellular Ca^2+^ (iCa^2+^) and maintenance of flatness in a steady-state epithelium. (A) Top view of a time-lapse movie of a MDCK confluent monolayer incubated with 5µM of Fura-2 dye showing intracellular calcium (iCa^2+^) levels. Selected sequential time frames are shown. (B, C) iCa^2+^ measurements (340nm/380nm) of 6 individual cells (different colored lines). (D) Model for iCa^2+^ regulation in a steady-state epithelium.

## Discussion

In this study, we employed several pharmacological agents to disrupt actin and myosin networks in a steady-state epithelium with the goal of understanding what forces are acting to maintain the flatness of epithelial sheets. Although there are individual caveats with each of the methods we used, when considered together the results hint at some interesting dynamics at play in maintaining a steady-state epithelium.

To quantify epithelial surface organization, we established a definition of flatness using confluent MDCK monolayers. Flatness was evaluated at both tissue, macro-, and cellular, micro-levels based on height variability across the epithelial surface. Confluent MDCK cells exhibited consistent uniform junctional organization marked by ZO-1, β-catenin, and phalloidin labeling. Using these measurements, we defined a flatness index based on the standard deviation of tissue and cellular height variation. Subconfluent MDCK cells, lacking full junctional organization, showed significantly higher variability, validating this metric as a measure of apical irregularity. Monolayers with standard deviations ≥1.0 μm (macro), or ≥0.4 μm (micro), were thus considered to have lost flatness. This framework provided a quantitative standard or threshold for assessing epithelial morphology and identifying disruptions in flatness following drug and genetic perturbations.

Epithelial height is regulated by both intracellular forces and ECM mechanics (Cai & Mostov, 2012; Janmey et al., 2020; Neelam et al., 2016). To examine whether the ECM topography influences apical geometry, MDCK cells were grown on patched laminin substrates that introduced uneven basement membrane contours. Two epithelial morphologies emerged: Type 1 epithelium on ‘steep’ laminin patches (ratio >2) formed wavy monolayers contiguous with the laminin contour, while Type 2 epithelium on ‘shallow’ patches (ratio <2) modulated individual cell heights relative to the laminin patch to maintain flatness. Type 1 epithelia exceeded the macro-height threshold, indicating a loss of apical flatness, whereas Type 2 epithelium remained within the defined standard. Micro-height was similar across all conditions, suggesting local height regulation despite topographical variation. These results demonstrate that ECM steepness, rather than composition, may dictate epithelial height organization: epithelial cells may sense the substrate slope and modulate their height to preserve overall epithelial architecture of the monolayer. Our finding of previously unknown epithelial tissue behavior in response to substrate steepness may provide new insights into epithelial cell morphology and plasticity and may broaden our understanding of cellular adaptation to complex microenvironments.

Interestingly, we did not observe any scenario in which only micro-height s.d. exceeded the flatness threshold while macro-height s.d. remained below it, indicating that micro-scale fluctuations alone do not drive loss of monolayer flatness. Instead, our cytoskeletal perturbation experiments showed that cytochalasin D, which disrupts actin, altered both macro- and micro-heights and led to loss of flatness, whereas nocodazole reduced micro-height without compromising overall flatness (Figure 3). These results highlight that loss of monolayer flatness is predominantly governed by changes in macro-height rather than micro-height. Monolayer flatness is an emergent property that may not directly reflect the behavior of individual cells but rather a tissue-level response.

To further investigate underlying cellular processes that may contribute to these responses, we pharmacologically inhibited actin and several myosin II-associated enzymes. These perturbations resulted in defected flatness, including increased micro-height and spherical “domes” of actin at the apical surface (Figures 3, 4). This phenotype was preserved following higher-specificity genetic perturbation with a mutant cell line, containing point mutations T696A and T853A to the MYPT1 regulatory subunit of MLCP. Interestingly, the transient transfection experiments also revealed that these mutations acted cell-autonomously, and exhibited no distinguishable effects on neighboring cells. Live imaging of iCa^2+^ fluorescence showed transient, asynchronous iCa^2+^ oscillations across the monolayer, suggesting a potential role of calcium, and further demonstrating cell autonomy in the process of maintaining epithelial flatness.

These data reveal several potential roles of myosin II in the maintenance of steady-state epithelial flatness. The first potential role is a detection mechanism to regulate cell height based on varying basal membrane topography/steepness. Our MDCK cells exhibited two distinct responses depending on the steepness of the laminin patch they were grown on. The exact mechanism of how MDCK cells sense ECM patch steepness, where the apical surface area is twice that of the basal surface, has not been previously reported, and remains to be explored.

Although myosin II has been known for its roles in the actomyosin contractile complex, it may also have non-contractile functions that are involved in flatness regulation of an epithelium. One potential role of myosin II in apical flatness is its influence on microvilli at the surface of MDCK cells. Pharmacological inhibition targeting myosin II ATPase, ROCK, and MLCK resulted in defective flatness, including increased micro-height and spherical “domes” of actin (Figure 4). Apical surface area enlargement and a dome-like spherical cap can be generated through microvilli retraction and buildup of apical actin (Chinowski et al., 2020; Ivanov et al., 2005; Jerdeva et al., 2005; Pietuch et al, 2013; Odenwald et al., 2018). Using TEM, we found a decrease in the number of microvilli in MDCK cells treated with ML-7 (sFigure 2), although the sample size analyzed is small. Future studies would benefit from increasing the sample size and examining the effects of blebbistatin or Y27632 treatment on microvilli number.

Our *MYPT1^T696A;T853A^*mutant cells displayed increased micro-height and spherical caps of actin (Figure 5), similar to the results of our myosin II inhibitor treatments. These data taken together support the model that myosin II may be functioning in multiple facets of flatness regulation at the apical surface of epithelial cells: as a component in the actomyosin-contractile complex, and in having a non-contractile role in microvilli architecture. The point mutations T696A and T853A are known to reduce phosphorylation of MYPT1 by ROCK, thus contributing to reduced MLCP inhibition and myosin II activity (Álvarez-Santos et al., 2020; Khasnis et al., 2014; Khromov et al., 2009; Koga & Ikebe, 2008). Importantly, no significant differences were observed in macro- or micro-height in GFP(-) cells adjacent to *MYPT1^T696A;T853A^* mutant cells, suggesting that maintenance of flatness is a cell-autonomous process. This finding is unexpected, as apical organization is often presumed to be tissue-coordinated, and it provides a mechanism by which epithelial integrity is preserved even when neighboring cells are perturbed.

Lastly, in a contractile/ tension model, actomyosin-generated mechanical force may create horizontal tension to keep micro- and macro-height consistent across the monolayer. Active myosin-actin interactions are triggered by local elevations of iCa^2+^ (Suzuki et al., 2017; Castillo et al., 1998; Sahu et al., 2017; Kuo & Ehrlich, 2015; Christodoulou & Skourides, 2015; Antunes et al., 2013). MDCK cells exhibit transient, asynchronous iCa^2+^ oscillations across the monolayer, allowing for iCa^2+^-dependent activation of MLCK-actomyosin complex and steady-state epithelial integrity while preventing harmful sustained elevations of iCa^2+^ (Figure 6) (Martin et al. 2009; Brodskiy & Zartman, 2018; Christodoulou & Skourides, 2015).

Additionally, since iCa^2+^ is a universal signaling molecule, we cannot exclude the possibility that indirect effects of iCa^2+^ oscillations contribute to the observed phenotypes as it is involved in many signaling pathways that regulate multiple essential functions (Kuo & Ehrlich, 2015; Brodskiy & Zartman, 2018; Moccia et al., 2023). For instance, while MLCK is directly activated by Ca^2+^, Ca^2+^ can also indirectly inhibit MLCP via PKC-mediated phosphorylation of CPI-17 (Woodsome et al., 2001), further influencing actomyosin dynamics. We analyzed iCa^2+^, but did not examine the potential effects of extracellular calcium. Future studies to investigate the role of extracellular calcium, and its contribution to the spatio-temporal properties of asynchronous Ca^2+^ transients of an epithelium to maintain flatness are needed.

This study sheds light into the dynamics of iCa^2+^-dependent MLCK and actomyosin complex in the physiology context of a steady-state epithelium. It is significant to understand regulators of apical epithelial cell morphology because loss of flatness has been correlated with tumor metastasis (Macara & McCaffrey, 2013; Jung et al., 2019; Hannezo et al., 2014; Eisenhoffer & Rosenblatt, 2012; Peglion & Etienne-Manneville 2024; Janiszewska et al., 2020; Ribatti et al., 2020; Coradini et al., 2011; Rodriguez-Boulan & Macara, 2014). By uncovering key mechanisms that govern apical morphology, this work contributes to our understanding of how disruptions in actomyosin dynamics and microvilli organization can drive changes in epithelial cytoarchitecture and function. Understanding how flatness is dynamically regulated may provide insight into epithelial resilience and failure during development, injury, and disease.

## Materials and Methods

### Cell preparation and culture

Madin Darby Canine Kidney (MDCK) cells (gift from Keith Mostov, UCSF) were cultured in DMEM complete media (Life technologies) supplemented with 5% FBS (Life Technologies), 1% penicillin-streptomycin in 10 cm^2^ cell culture plates (Corning) and/or transwell cell culture inserts (Corning).

### Transient transfection of mutant MDCK cell lines

*MYPT1^T696A;T853A^* mutant MDCK lines were generated by plasmid transfection using the Neon electroporation system (Invitrogen). The insert was cloned using Gibson assembly into the pK.myc-MYPT1 plasmid (#24101, Addgene). Mutations were introduced in the plasmid using the In-Fusion Cloning kit (TakaraBio) primer design approach. Primer pairs covering the cDNA sequences were designed: 5’ GTC TAG AAG ATC AGC ACA GGG AGT AAC ATT GAC T 3’, and 5’ AGT CAA TGT TAC TCC CTG TGC TGA TCT TCT AGA C 3’ for *MYPT1 T695A point mutation*; 5’ GAG AAA AGG AGA TCC GCA GGA GTT TCA TTT TGG ACA CA 3’, and 5’ TGT GTC CAA AAT GAA ACT CCT GCG GAT CTC CTT TTC TC 3’ for *MYPT1 T852A point mutation*.

The constructed vectors were then sequenced (Quintara Biosciences), and used for transfection.

### Transepithelial electrical resistance (TEER)

MDCKs were harvested from 10 cm^2^ cell culture plates at a confluency of 70-80%, and were seeded at a density of 1–5 × 10^5^ cells per insert in the Transwell membrane apparatuses (Corning) to desired confluence. TEER measurements were taken 24 hr after seeding, at each 1 hr time point in a Biosafety cabinet at room temperature (RT) using an Epithelial Volt/Ohm meter (EVOM) (World Precision Instruments). The background resistance was taken from an insert with media only, and subtracted for each TEER reading, and multiplied by the surface area of the filter (1.12 cm^2^).

### Drug treatments

MDCK cells were treated with 10µM of cytochalasin D (C8273, Sigma), or 5µM of withaferin A (W4394, Sigma), or 16.5µM of nocodazole (M1404, Sigma), 30µM blebbistatin (B0560, Sigma) or 50µM Y27632 (sc-281642; Santa Cruz Biotechnology), 100µM ML-7 (ab120848; Abcam), dissolved in DMSO for 2 hr in an incubator at 37°C, 5% CO_2_ before processing. Experiments were completed on three individual MDCK monolayer samples for each drug treatment. Fifteen cells were measured from each monolayer sample for a total of 45 cells per treatment.

### Immunofluorescence

Transwell membranes were transferred to a 12-well plate containing cold phosphate-buffered saline (PBS). The permeable Transwell membranes were then fixed with 4% paraformaldehyde (PFA) for 30-60 min on ice, rinsed 3X with cold PBS, and incubated with permeabilization solution (PFS) (0.7% fish skin gelatin, 0.025% saponin dissolved in PBS) for 1 hr at RT. Membranes were then cut from the original Transwell apparatus using a scalpel. Membranes were then incubated with antibodies to mouse ZO-1 (1:50) (Abcam), rabbit β-catenin (1:1000) (ab6302, Abcam), stained with Alexa Fluor 647 phalloidin (1:500) (A22287, Invitrogen), and DAPI (1:1000) (Invitrogen). Secondary antibodies using rabbit Alexa Fluor 488 (1:100) (Invitrogen), Alexa Fluor 594 (Invitrogen) were then applied. Following antibody staining, Transwell membranes were then mounted on glass slides (FisherScientific) using Prolong Gold Antifade (Invitrogen), and coverslipped (FisherScientific). Rhodamine-laminin (#LMN01-A, Cytoskeleton, Inc) was applied to Transwell membranes. Following antibody staining, Transwell membranes were then mounted on glass slides (FisherScientific) using Prolong Gold Antifade (Invitrogen), and coverslipped (FisherScientific).

### Image capture and Acquisition

Fixed MDCK cells were imaged using a confocal microscope (TCS SP2; Leica) using a 40X, and 63X oil immersion objective as previously described (Hua and Mikawa, 2018). Images were reconstructed from confocal optical sections using Imaris software (Bitplane). Brightness and color were adjusted using Photoshop (Adobe Systems).

### Transmission electron microscopy (TEM)

MDCK cells were seeded in Transwell filters, and transwell filters cut into small pieces and put into embedding plastic molds. After overnight adherence, the cells were either treated with DMSO (control) or 100µM ML-7. The morphology of the MDCK cells were visualized with TEM as described in Heckman et al., (2007).

### Quantification of cell and apical height analysis

MDCK monolayers were reconstructed from confocal optical sections, and analyzed using Imaris software (Bitplane). Measurements were conducted using the Imaris measurement tool function on an orthogonal plane from the 3D reconstructed optical sections (sFig 1 C). For immunofluorescence analysis, fluorescent images were thresholded by highlighting the brightest 80% of the images for all channels. Phalloidin staining was used to determine the maximum height of the cell’s apex (sFig 1 C). Micro-height was measured as the vertical distance from tricellular junction midpoint to the apex of phalloidin staining (sFig 1 C). Macro-height was measured as the vertical distance from the midpoint to the cell base (sFig 1 C).

Plots were generated in Graphpad Prism10 to present height measurements, treated vs untreated cells, GFP(-), GFP(+) cells for the various experiments, and ratio of apical/basal area of laminin patches in MDCK culture conditions. Quantification of ECM bumps was completed using Imaris (Bitplane).

### Statistical analysis

Confluent and subconfluent measurements (Figure 1) and cell height measurements grown on laminin patches (Figure 2) were statistically compared using either a parametric or non-parametric test, contingent upon Levene’s test for unequal variance between groups. Equal variance prompts a parametric student’s t-test, while unequal variance prompts a non-parametric Mann-Whitney U-test. Drug-treated samples (Untreated, cytochalasin D, withaferin A, nocodazole (Figure 3)), (blebbistatin, Y27632, ML-7 (Figure 4)), and transfected samples (*MYPT1^T696A;T853A^*; Figure 5) were statistically compared using either a One-way ANOVA or Kruskal-Wallis test, contingent upon Levene’s test for unequal variance between groups. Equal variance prompts a parametric One-Way ANOVA, while unequal variance prompts a non-parametric Kruskal-Wallis test. On Figures 3 and 4, post-hoc tests (Dunnett’s test if a One Way ANOVA or Dunn’s test if a Kruskal-Wallis test) were performed, allowing multiple comparisons of the treated groups to the DMSO control group.

### Calcium imaging

MDCK cells were grown on 3.5 cm glass dishes (Ibidi) and incubated with 2µM Fura-2 AM in DMSO at 37°C for 30 minutes. Cells were then washed with 1X Hank’s Balanced Salt Solution (HHBS). Individual cells were excited at 340 nm and 380 nm with emission at 510nm as a ratiometric indicator for intracellular Ca^2+^concentration measurements. Ex/Em: 340/510 nm can be used to measure Ca^2+^-bound Fura-2, and Ex/Em: 380/510 nm can be used to measure Ca^2+^-free Fura-2, indicating high and low concentrations of Ca^2+^ (Ion Biosciences). Imaging was done at 300ms intervals with no delay for 20 minutes. Imaging was captured using an inverted Nikon Ti CSU-X1 Spinning Disk confocal microscope supported by a Photometrics Prime 95B sCMOS camera, and an OkoLab cage incubator with CO_2_ and humidity control with a Plan Apo 40X Objective using Nikon Elements Advance Research software.

## Acknowledgements

We thank Dr. Keith Mostov for providing the MDCK cells; T.M. and L.L.H laboratory members for their comments and suggestions: Sara Venters, Michael Bressan; L.L.H laboratory member Gabriel Quintero Plancarte for his guidance and insight; SSU Biology Faculty: Dr. Daniel Crocker for his statistics expertise and Dr. Joseph Lin for his helpful feedback; SSU Women’s Basketball Coaches: Des Abeyta and Trevor Johnson; and the Herrmann Family: Ryan, Susan, Jack, Reese, and Blue for their encouragement and support. L.L.H dedicates this work to the memory of her daughter, Josephine Jereb. We apologize to the colleagues whose work we could not cite due to space limitations. This work was supported in part by National Institutes of Health (NIH) Awards (#R01HL148125 and #R01HL153736 to T. M. and #R16GM153517, to L.L.H), National Science Foundation (NSF RUI Award #2027746, to L.L.H.), California State University Program for Education and Research in Biotechnology (CSUBIOTECH Research Development Grant to L.L.H.), Smith Family Funds to T.M, SSU NCAA Basketball Athletic Scholarships to E.G.H.

## Supplementary Figure Captions

**sFigure 1. Characterization and quantification of MDCK cells.**

(A) TEER measurements for MDCK cells grown in culture. MDCK monolayers displayed the highest electrical resistance of 279.6 Ω*cm^2^ s.d. 33.23 at 45 hours over a 3-day measurement period of the same monolayer. (B) Schematic of individual MDCK cell selection from a monolayer. (B’) Selected cell in (B) with measurement points (yellow). Measurements were taken by identifying the midpoint (yellow dot) of the apical actin belt using ZO-1 staining (red) and drawing a line (yellow horizontal line). Macro-height was determined by the distance from the midpoint (yellow dot) to the cell base (vertical yellow line). (B”) Selected apical domain in (B’). Micro-height was measured as the distance from the midpoint to the peak of phalloidin staining, h, (grey). (C) Steps taken to quantification of macro- and micro-height: 1) Top view of stacked confocal sections of a confluent MDCK monolayer. 2) Side view of the monolayer after a 90° clockwise rotation along the x-axis. 3) X-Z orthogonal plane showing a cross-section of the monolayer for side-view cell measurements. 4) Select individual cell for quantification. 5) Zoom in on the selected cell. 6) Measurement of the apical horizontal line connecting tight junctions via ZO-1. 7) Micro-height measurement as described in (B”). 8) Macro-height measurement as described in (B’). Fluorescent images were thresholded to highlight the top 80% brightness across all channels. (D) Top and side views of stacked confocal optical sections a MDCK subconfluent culture immunostained with ZO-1 (red), β-catenin (green), and filamentous actin dye, phalloidin (grey), and DNA DAPI (blue) counterstain. (Scale bars: 2, 10µm)

**sFigure 2. Transmission electron microscopy (TEM) reveals decreased number of microvilli in ML-7 treated MDCK cells.**

(A)Transmission electron microscopy (TEM) for MDCK cells treated with DMSO. (B) As in (A) but treated with 100 µM ML-7. (C) Quantification of microvilli number of DMSO with an average of 35.88 ± 6.13 s.d. and ML-7 treated with an average of 6.75 ± 3.54 s.d. number of microvilli (n=8 cells for each treatment) (p<0.0001, t=11.64). Treatments were statistically compared with a Student’s t-test. Note: ML-7 treated MDCK cells showed a decrease of microvilli number. (Scale bars: 2 µm)

**sMovie1. Live cell imaging of intracellular Ca^2+^ (iCa^2+^) and maintenance of apical flatness in steady state epithelium.**

Live imaging video of an MDCK monolayer acquired over 30 minutes at 300 ms intervals stained with Fura-2 calcium dye. High and low [iCa^2+^] are designated in red and green, respectively. Note: iCa^2+^ concentrations oscillate asynchronously among individual MDCK cells across the monolayer. (Scale bars: 2 µm)

## References

Álvarez-Santos, M. D., Álvarez-González, M., Estrada-Soto, S., Bazán-Perkins, B. (2020). Regulation of Myosin Light-Chain Phosphatase Activity to Generate Airway Smooth Muscle Hypercontractility. Sec. Respiratory Physiology and Pathophysiology, 11:701. 10.3389/fphys.2020.00701

Antunes, M., Pereira, T., Cordeiro, J. V., Almeida, L., & Jacinto, A. (2013). Coordinated waves of actomyosin flow and apical cell constriction immediately after wounding. The Journal of cell biology, 202(2), 365–379. 10.1083/jcb.201211039

Bagur, R., & Hajnóczky, G. (2017). Intracellular Ca2+ Sensing: Its Role in Calcium Homeostasis and Signaling. Molecular cell, 66(6), 780–788. 10.1016/j.molcel.2017.05.028

Banjac, I., Maimets, M., Jensen, K. B. (2023). Maintenance of high-turnover tissues during and beyond homeostasis. Cell Stem Cell, 30(4), 348–361. 10.1016/j.stem.2023.03.008

Barlan, K., & Gelfand, V. I. (2017). Microtubule-based transport and the distribution, tethering, and organization of organelles. Cold Spring Harbor perspectives in biology, 9(5), a025817. 10.1101/cshperspect.a025817

Barreto-Chang, O. L., & Dolmetsch, R. E. (2009). Calcium imaging of cortical neurons using Fura-2 AM. Journal of visualized experiments : JoVE, (23), 1067. 10.3791/1067

Bers, D. M. (2008). Calcium cycling and signaling in cardiac myocytes. Annual Review of Physiology, 70. 10.1146/annurev.physiol.70.113006.100455

Bresnick A. R. (1999). Molecular mechanisms of nonmuscle myosin-II regulation. Current opinion in cell biology, 11(1), 26–33. 10.1016/s0955-0674(99)80004-0

Cai, L., & Mostov, K. E. (2012). Cell height: Tao rising. The Journal of cell biology, 199(7), 1023–1024. 10.1083/jcb.201211015

Castillo, A. M., Lagunes, R., Urban, M., Frixione, E., & Meza, I. (1998). Myosin II-actin interaction in MDCK cells: role in cell shape changes in response to Ca2+ variations. Journal of muscle research and cell motility, 19(5), 557–574. 10.1023/a:1005316711538

Chinowsky, C. R., Pinette, J. A., Meenderink, L. M., Lau, K. S., and Tyska, M. J. (2020). Nonmuscle myosin-2 contractility-dependent actin turnover limits the length of epithelial microvilli. Molecular Biology of the Cell, 31(25). 10.1091/mbc.E20-09-0582

Christodoulou, N., & Skourides, P. A. (2015). Cell-Autonomous Ca(2+) Flashes Elicit Pulsed Contractions of an Apical Actin Network to Drive Apical Constriction during Neural Tube Closure. Cell reports, 13(10), 2189–2202. 10.1016/j.celrep.2015.11.017

Chugh P., Paluch E. K. (2018). The actin cortex at a glance. J Cell Sci, 131(14), jcs186254. doi: 10.1242/jcs.186254.

Clarke, D. N., & Martin, A. C. (2021). Actin-based force generation and cell adhesion in tissue morphogenesis. Current biology : CB, 31(10), R667–R680. 10.1016/j.cub.2021.03.031

Coradini, D., Casarsa, C. & Oriana, S. (2011). Epithelial cell polarity and tumorigenesis: new perspectives for cancer detection and treatment. Acta Pharmacol Sin, 32, 552–564. 10.1038/aps.2011.20

Cui, Wj., Liu, Y., Zhou, Xl. et al. (2010). Myosin light chain kinase is responsible for high proliferative ability of breast cancer cells via anti-apoptosis involving p38 pathway. Acta Pharmacol Sin, 31, 725–732. 10.1038/aps.2010.56

Diaz-de-la-Loza, M., Ray, R. P., Ganguly, P. S., Alt, S., Davis, J. R., Hoppe, A., Tapon, N., Salbreux, G., Thompson, B. J. (2018). Apical and basal matrix remodeling control epithelial morphogenesis. Developmental Cell, 46(1), 23–39.e5. 10.1016/j.devcel.2018.06.006

Dukes, J. D., Whitley, P., & Chalmers, A. D. (2011). The MDCK variety pack: choosing the right strain. BMC Cell Biol, 12, 43. 10.1186/1471-2121-12-43

Ebrahim, S., Fujita, T., Millis, B. A., Kozin, E., Ma, X., Kawamoto, S., Baird, M. A., Davidson, M., Yonemura, S., Hisa, Y., Conti, M. A., Adelstein, R. S., Sakaguchi, H., & Kachar, B. (2013). NMII forms a contractile transcellular sarcomeric network to regulate apical cell junctions and tissue geometry. Current biology : CB, 23(8), 731–736. 10.1016/j.cub.2013.03.039

Eisenhoffer, G. T., & Rosenblatt, J. (2012). Bringing balance by force: live cell extrusion controls epithelial cell numbers. Trends in Cell Biology, 23(4) 185–192. 10.1016/j.tcb.2012.11.006

Fanning, A. S., Van Itallie, C. M., & Anderson, J. M. (2012). Zonula occludens-1 and -2 regulate apical cell structure and the zonula adherens cytoskeleton in polarized epithelia. Molecular biology of the cell, 23(4), 577–590. 10.1091/mbc.E11-09-0791

Fedi, A., Vitale, C., Ponschin, G., Ayehunie, S., Fato, M., Scaglione, S. (2021). In vitro models replicating the human intestinal epithelium for absorption and metabolism studies: A systematic review. Journal of controlled release, 335, 247–268. 10.1016/j.jconrel.2021.05.028.

Frantz, C., Stewart, K. M., & Weaver, V. M. (2010). The extracellular matrix at a glance. Journal of cell science, 123(Pt 24), 4195–4200. 10.1242/jcs.023820

Gaeta, I. M., Meenderink, L. M., Postema, M. M., Cencer, C. S., & Tyska, M. J. (2021). Direct visualization of epithelial microvilli biogenesis. Current Biology, 31(12), 2561–2575.e6, ISSN 0960-9822. 10.1016/j.cub.2021.04.012

Garrido-Casado, M., Asensio-Juárez, G., Talayero, V. C., & Vicente-Manzanares, M. (2024). Engines of change: Nonmuscle myosin II in mechanobiology. Current Opinion in Cell Biology, 87, 102344, ISSN 0955-0674. 10.1016/j.ceb.2024.102344

Goeckeler, Z. M., Bridgman, P. C., & Wysolmerski, R. B. (2008). Nonmuscle myosin II is responsible for maintaining endothelial cell basal tone and stress fiber integrity. American journal of physiology. Cell physiology, 295(4), C994–C1006. 10.1152/ajpcell.00318.2008

Hannezo, E., Prost, J., & Joanny, J. F. (2014). Theory of epithelial sheet morphology in three dimensions. Proceedings of the National Academy of Sciences of the United States of America, 111(1), 27–32. 10.1073/pnas.1312076111

Hathaway, D.R., & Adelstein, R. S. (1979). Human platelet myosin light chain kinase requires the calcium-binding protein calmodulin for activity., Proc. Natl. Acad. Sci. U.S.A. 76(4) 1653–1657, 10.1073/pnas.76.4.1653

Heckman, C., Kanagasundaram, S., Cayer, M., Paige, J. (2007). Preparation of cultured cells for scanning electron microscope. Protocol Exchange; doi: 10.1038/nprot.2007.504

Hirokawa, N., Tilney, L. G., Fujiwara, K., & Heuser, J. E. (1982). Organization of actin, myosin, and intermediate filaments in the brush border of intestinal epithelial cells. The Journal of cell biology, 94(2), 425–443. 10.1083/jcb.94.2.425

Hua, L. L., & Mikawa, T. (2018). Mitotic antipairing of homologous and sex chromosomes via spatial restriction of two haploid sets. Proceedings of the National Academy of Sciences of the United States of America, 115(52), E12235–E12244. 10.1073/pnas.1809583115

Huang, L., Peng, Y., Tao, X., Ding, X., Li, R., Jiang, Y., & Zuo, W. (2022). Microtubule Organization Is Essential for Maintaining Cellular Morphology and Function. Oxidative medicine and cellular longevity, 1623181. 10.1155/2022/1623181

Ivanov, A. I., Hunt, D., Utech, M., Nusrat, A., & Parkos, C. A. (2005). Differential roles for actin polymerization and a myosin II motor in assembly of the epithelial apical junctional complex. Molecular biology of the cell, 16(6), 2636–2650. 10.1091/mbc.e05-01-0043

Janiszewska, M., Primi, M. C., & Izard, T. (2020). Cell adhesion in cancer: Beyond the migration of single cells. Journal of Biological Chemistry, 295(8), 2495–2505, ISSN 0021-9258. 10.1074/jbc.REV119.007759.

Janmey, P. A., Fletcher, D. A., & Reinhart-King, C. A. (2020). Stiffness Sensing by Cells. Physiological reviews, 100(2), 695–724. 10.1152/physrev.00013.2019

Jerdeva, G. V., Wu, K., Yarber, F. A., Rhodes, C. J., Kalman, D., Schechter, J. E., & Hamm-Alvarez, S. F. (2005). Actin and non-muscle myosin II facilitate apical exocytosis of tear proteins in rabbit lacrimal acinar epithelial cells. Journal of cell science, 118(Pt 20), 4797–4812. 10.1242/jcs.02573

Jung, H. Y., Fattet, L., Tsai, J. H., Kajimoto, T., Chang, Q., Newton, A. C., & Yang, J. (2019). Apical-basal polarity inhibits epithelial-mesenchymal transition and tumour metastasis by PAR-complex-mediated SNAI1 degradation. Nature cell biology, 21(3), 359–371. 10.1038/s41556-019-0291-8

Kassianidou, E., Hughes, J. H., & Kumar, S. (2017). Activation of ROCK and MLCK tunes regional stress fiber formation and mechanics via preferential myosin light chain phosphorylation. Molecular biology of the cell, 28(26), 3832–3843. 10.1091/mbc.E17-06-0401

Khalilgharibi, N., & Mao, Y. (2021). To form and function: on the role of basement membrane mechanics in tissue development, homeostasis and disease. Open biology, 11(2), 200360. 10.1098/rsob.200360

Khasnis, M., Nakatomi, A., Gumpper, K., & Eto, M. (2014). Reconstituted human myosin light chain phosphatase reveals distinct roles of two inhibitory phosphorylation sites of the regulatory subunit, MYPT1. Biochemistry, 53, 2701–2709. 10.1021/bi5001728

Khromov, A., Choudhury, N., Stevenson, A. S., Somlyo, A. V., & Eto, M. (2009). Phosphorylation-dependent autoinhibition of myosin light chain phosphatase accounts for Ca^2+^ sensitization force of smooth muscle contraction. Journal of Biological Chemistry, 284(32) 21569–21579, ISSN 0021-9258. 10.1074/jbc.M109.019729

Klingner, C., Cherian, A.V., Fels, J., Diesinger, P. M., Aufschnaiter, R., Maghelli, N., Keil, T., Beck, G., Tolić-Nørrelykke, I. M., Bathe, M., Wedlich-Soldner, R. (2014). Isotropic actomyosin dynamics promote organization of the apical cell cortex in epithelial cells. J Cell Biol, 207(1), 107–121. 10.1083/jcb.201402037

Kondo, T., & Hayashi, S. (2015). Mechanisms of cell height changes that mediate epithelial invagination. Develop. Growth Differ., 57, 313–323. 10.1111/dgd.12224

Kovács, M., Tóth, J., Hetényi, C., Málnási-Csizmadia, A., & Sellers, J. R. (2004). Mechanism of blebbistatin inhibition of myosin II. The Journal of biological chemistry, 279(34), 35557–35563. 10.1074/jbc.M405319200

Kozyrina, A. N., Piskova, T., Di Russo, J. (2020). Mechanobiology of epithelia from the perspective of extracellular matrix heterogeneity. Front. Bioeng. Biotechnol, 8. 10.3389/fbioe.2020.596599

Kuo, I. Y., & Ehrlich, B. E. (2015). Signaling in muscle contraction. Cold Spring Harbor perspectives in biology, 7(2), a006023. 10.1101/cshperspect.a006023

Kuo, W. T., Odenwald, M. A., Turner, J. R., Zuo, L. (2022). Tight junction proteins occludin and ZO-1 as regulators of epithelial proliferation and survival. Ann N Y Acad Sci, 1514(1), 21–33. doi: 10.1111/nyas.14798.

Lee, L. W., Lee, G. H., Su, I. H., Lu, C. H., Lin, K. H., Wen, F. L. & Tang, M. J. (2025). Mechanobiological mechanism of cyclic stretch-induced cell columnarization. Cell Reports, 44(5), 115662, ISSN 2211-1247, 10.1016/j.celrep.2025.115662

Lemke, S. B., Nelson, C. M. (2021). Dynamic changes in epithelial cell packing during tissue morphogenesis. Current Biology, 31(18), R1098–R1110. 10.1016/j.cub.2021.07.078

Leng, S., Pignatti, E., Khetani, R.S. et al. (2020). β-Catenin and FGFR2 regulate postnatal rosette-based adrenocortical morphogenesis. Nat Commun 11, 1680. 10.1038/s41467-020-15332-7

Macara, I. G., & McCaffrey, L. (2013). Cell polarity in morphogenesis and metastasis. Philosophical transactions of the Royal Society of London. Series B, Biological sciences, 368(1629), 20130012. 10.1098/rstb.2013.0012

Marivin, A., Ho, R. X., & Garcia-Marcos, M. (2022). DAPLE orchestrates apical actomyosin assembly from junctional polarity complexes. The Journal of cell biology, 221(5), e202111002. 10.1083/jcb.202111002

Martin, A. C, Goldstein, B. (2014). Apical constriction: themes and variations on a cellular mechanism driving morphogenesis. Development, 141(10), 1987–98. doi: 10.1242/dev.102228

Martin, A. C., Kaschube, M., & Wieschaus, E. F. (2009). Pulsed contractions of an actin-myosin network drive apical constriction. Nature, 457(7228), 495–499. 10.1038/nature07522

Martin-Belmonte, F., & Mostov, K. (2008). Regulation of cell polarity during epithelial morphogenesis, Current Opinion in Cell Biology, 20(2), 227–234, 10.1016/j.ceb.2008.01.001

Meng, W., & Takeichi, M. (2009). Adherens junction: molecular architecture and regulation. Cold Spring Harbor perspectives in biology, 1(6), a002899. 10.1101/cshperspect.a002899

McNeil, E., Capaldo, C. T., & Macara, I. G. (2006). Zonula occludens-1 function in the assembly of tight junctions in Madin-Darby canine kidney epithelial cells. Molecular biology of the cell, 17(4), 1922–1932. 10.1091/mbc.e05-07-0650

Misfeldt, D. S., Hamamoto, S. T., & Pitelka, D. R. (1976). Transepithelial transport in cell culture. Proceedings of the National Academy of Sciences of the United States of America, 73(4), 1212–1216. 10.1073/pnas.73.4.1212

Moccia, F., Fiorio, A., Lim, D., Lodola, F., & Gerbino, A. (2023). Intracellular Ca^2+^ signalling: unexpected new roles for the usual suspect. Front. Physiol., 14. 10.3389/fphys.2023.1210085

Mooseker, M. S., & Tilney, L. G. (1975). Organization of an actin filament-membrane complex. Filament polarity and membrane attachment in the microvilli of intestinal epithelial cells. J Cell Biol, 67(3): 725–743. 10.1083/jcb.67.3.725

Mortensen, K., Larsson, LI. (2003). Effects of cytochalasin D on the actin cytoskeleton: association of neoformed actin aggregates with proteins involved in signaling and endocytosis. *CMLS*, Cell. Mol. Life Sci, 60, 1007–1012. 10.1007/s00018-003-3022-x

Muncie, J. M., & Weaver, V. M. (2018). The physical and biochemical properties of the extracellular matrix regulate cell fate. Current topics in developmental biology, 130, 1–37. 10.1016/bs.ctdb.2018.02.002

Neelam, S., Hayes, P., Zhang, Q., Dickinson, R. B., & Lele, T. P. (2016). Vertical uniformity of cells and nuclei in epithelial monolayers. Sci Rep, 6, 19689. 10.1038/srep19689

Odenwald, M. A., Choi, W., Kuo, W. T., Singh, G., Sailer, A., Wang, Y., Shen, L., Fanning, A. S., & Turner, J. R. (2018). The scaffolding protein ZO-1 coordinates actomyosin and epithelial apical specializations in vitro and in vivo. The Journal of biological chemistry, 293(45), 17317–17335. 10.1074/jbc.RA118.003908

Oelz, D. B., del Castillo, U., Gelfand, V. I., & Mogilner, A. (2018). Microtubule dynamics, kinesin-1 sliding, and dynein action drive growth of cell processes. Biophysical Journal, 115(8), 1614–1624. 10.1016/j.bpj.2018.08.046

Paredes, M. R., Etzler, J.C., Watts, L.T., Zheng, W., & Lechleiter, J. D. (2008). Chemical calcium indicators. Methods, 46(3), 143–151. 10.1016/j.ymeth.2008.09.025

Peglion, F., & Etienne-Manneville, S. (2024). Cell polarity changes in cancer initiation and progression. The Journal of cell biology, 223(1), e202308069. 10.1083/jcb.202308069

Perez-Tirado, A., Unkelbach, U., Oswald, T. A., Rheinlaender, J., Schäffer, T. E., Mukenhirn, M., Honigmann, A., Janshoff, A. (2025). Differences in apical and basal mechanics regulate compliance of curved epithelia. Cell Reports Physical Science, 6(3), 102485. 10.1016/j.xcrp.2025.102485

Pietuch, A., Brückner, B. R., & Janshoff, A. (2013). Membrane tension homeostasis of epithelial cells through surface area regulation in response to osmotic stress. Biochimica et Biophysica Acta (BBA) - Molecular Cell Research, 1833(3), 712–722, 10.1016/j.bbamcr.2012.11.006

Pollard, T. D., & Cooper, J. A. (2009). Actin, a central player in cell shape and movement. *Science (New York*, N.Y.), 326(5957), 1208–1212. 10.1126/science.1175862

Quintin, S., Gally, C., & Labouesse, M. (2008). Epithelial morphogenesis in embryos: asymmetries, motors and brakes. Trends in genetics : TIG, 24(5), 221–230. 10.1016/j.tig.2008.02.005

Ranie, S. N., & White, M. D. (2025). Apical constriction in morphogenesis: From actomyosin architecture to regulatory networks. Current Opinion in Cell Biology, 95, 102562, ISSN 0955-0674. 10.1016/j.ceb.2025.102562

Ribatti, D., Tamma, R., & Annese, T. (2020). Epithelial-Mesenchymal Transition in Cancer: A Historical Overview. Translational oncology, 13(6), 100773. 10.1016/j.tranon.2020.100773

Rodriguez-Boulan, E., & Macara, I. (2014) Organization and execution of the epithelial polarity programme. Nat Rev Mol Cell Biol, 15, 225–242. 10.1038/nrm3775

Rozario, T., & DeSimone, D. W. (2010). The extracellular matrix in development and morphogenesis: a dynamic view. Developmental biology, 341(1), 126–140. 10.1016/j.ydbio.2009.10.026

Sahu, S. U., Visetsouk, M. R., Garde, R. J., Hennes, L., Kwas, C., & Gutzman, J. H. (2017). Calcium signals drive cell shape changes during zebrafish midbrain-hindbrain boundary formation. Molecular biology of the cell, 28(7), 875–882. 10.1091/mbc.E16-08-0561

Schliwa M. (1982). Action of cytochalasin D on cytoskeletal networks. The Journal of cell biology, 92(1), 79–91. 10.1083/jcb.92.1.79

Sharkova, M., Chow, E., Erickson, T., & Hocking, J. C. (2023). The morphological and functional diversity of apical microvilli. Journal of anatomy, 242(3), 327–353. 10.1111/joa.13781

Shibata, K., Sakai, H., Huang, Q., Kamata, H., Chiba, Y., Misawa, M., Ikebe, R., & Ikebe, M. (2015). Rac1 regulates myosin II phosphorylation through regulation of myosin light chain phosphatase. Journal of cellular physiology, 230(6), 1352–1364. 10.1002/jcp.24878

Srinivasan, B., Kolli, A. R., Esch, M. B., Abaci, H. E., Shuler, M. L., & Hickman, J. J. (2015). TEER measurement techniques for in vitro barrier model systems. Journal of laboratory automation, 20(2), 107–126. 10.1177/2211068214561025

Suzuki, M., Sato, M., Koyama, H., Hara, Y., Hayashi, K., Yasue, N., Imamura, H., Fujimori, T., Nagai, T., Campbell, R. E., & Ueno, N. (2017). Distinct intracellular Ca^2+^ dynamics regulate apical constriction and differentially contribute to neural tube closure. *Development (Cambridge*, England), 144(7), 1307–1316. 10.1242/dev.141952

Tai, K., Cockburn, K., & Greco, V. (2019). Flexibility sustains epithelial tissue homeostasis. Current opinion in cell biology, 60, 84–91. 10.1016/j.ceb.2019.04.009

Tang V. (2017). Cell-cell adhesion interface: rise of the lateral membrane. F1000Research, 6, 276. 10.12688/f1000research.10680.1

Töpfer U. (2023). Basement membrane dynamics and mechanics in tissue morphogenesis. Biology open, 12(8), bio059980. 10.1242/bio.059980

Totsukawa, G., Yamakita, Y., Yamashiro, S., Hartshorne, D. J., Sasaki, Y., & Matsumura, F. (2000). Distinct Roles of Rock (Rho-Kinase) and Mlck in Spatial Regulation of Mlc Phosphorylation for Assembly of Stress Fibers and Focal Adhesions in 3t3 Fibroblasts. J Cell Biol, 150(4), 797–806. 10.1083/jcb.150.4.797

Vasquez, C. G., Tworoger, M., & Martin, A. C. (2014). Dynamic myosin phosphorylation regulates contractile pulses and tissue integrity during epithelial morphogenesis. The Journal of cell biology, 206(3), 435–450. 10.1083/jcb.201402004

Watanabe, T., Hosoya, H., & Yonemura, S. (2007). Regulation of myosin II dynamics by phosphorylation and dephosphorylation of its light chain in epithelial cells. Molecular biology of the cell, 18(2), 605–616. 10.1091/mbc.e06-07-0590.

Wells, C. L., van de Westerlo, E. M., Jechorek, R. P., Haines, H. M., & Erlandsen, S. L. (1998). Cytochalasin-induced actin disruption of polarized enterocytes can augment internalization of bacteria. Infection and immunity, 66(6), 2410–2419. 10.1128/IAI.66.6.2410-2419.1998

Wells, E. K., Yarborough, O., 3rd, Lifton, R. P., Cantley, L. G., & Caplan, M. J. (2013). Epithelial morphogenesis of MDCK cells in three-dimensional collagen culture is modulated by interleukin-8. American journal of physiology. Cell physiology, 304(10), C966–C975. 10.1152/ajpcell.00261.2012

Woodsome, T. P., Eto, M., Everett, A., Brautigan, D. L., & Kitazawa, T. (2001). Expression of CPI-17 and myosin phosphatase correlates with Ca(2+) sensitivity of protein kinase C-induced contraction in rabbit smooth muscle. The Journal of physiology, 535(Pt 2), 553–564. 10.1111/j.1469-7793.2001.t01-1-00553.x

Yerna, M. J., Dabrowska, R., Hartshorne, D. J., & Goldman, R. D. (1979). Calcium-sensitive regulation of actin-myosin interactions in baby hamster kidney (BHK-21) cells., Proc. Natl. Acad. Sci. U.S.A. 76 (1) 184–188, 10.1073/pnas.76.1.184

